# Mortality without springing a leak: Locust gut epithelia do not become more permeable to fluorescent dextran and bacteria in the cold

**DOI:** 10.1101/2022.09.21.508851

**Authors:** Mahmoud I. El-Saadi, Kaylen Brzezinski, Aaron Hinz, Laura Phillips, Alex Wong, Lucie Gerber, Johannes Overgaard, Heath A. MacMillan

**Affiliations:** Department of Biology, Carleton University, Ottawa, Canada, K1S 5B6; Department of Bioscience, University of Oslo, Oslo, Norway; Department of Biology, Aarhus University, Aarhus, Denmark

**Keywords:** Chilling injury, gut bacteria, microbiome, immunity, paracellular leak, dextran

## Abstract

The insect gut, which plays a role in ion and water balance, has been shown to leak solutes in the cold. Cold stress can also activate insect immune systems, but it is unknown if the leak of the gut microbiome is a possible immune trigger in the cold. We developed a novel feeding protocol to load the gut of locusts (*Locusta migratoria*) with fluorescent bacteria before exposing them to -2°C for up to 48 h. No bacteria were recovered from the hemolymph of cold-exposed locusts, regardless of exposure duration. To examine this further, we used an *ex vivo* gut sac preparation to re-test cold-induced fluorescent FITC-dextran leak across the gut and found no increased rate of leak. These results question not only the validity of FITC-dextran as a marker of paracellular barrier permeability in the gut, but also to what extent the insect gut becomes leaky in the cold.

## Introduction

The majority of insects are chill-susceptible, meaning they suffer negative effects of chilling (chilling injuries) at low temperatures well above the freezing point of their body fluids (Overgaard and MacMillan, 2017). As temperatures drop below an insect’s critical thermal minimum (CTmin), they lose coordinated motor control. Continued cold exposure eventually leads to the onset of chill-coma, a state characterized by a complete but reversible paralysis (Andersen et al., 2015; Hazell and Bale, 2011; MacMillan and Sinclair, 2011; Rodgers et al., 2010). Prolonged exposure to low temperatures leads to tissue damage in the insect, and these chilling injuries can accumulate and increase in severity if temperatures remain. Chilling injuries are thought to be largely driven by cell death resulting from a loss of ion homeostasis (Andersen et al., 2017a; Bayley et al., 2018; Carrington et al., 2020; Koštál et al., 2004; MacMillan and Sinclair, 2011b; Overgaard et al., 2021). Water and ion balance play a key role in maintaining neuromuscular function, but at low temperatures, active transport rates of solutes are slowed to a point where they cannot counterbalance the passive leak of solutes and water (Overgaard et al., 2021).

The insect gut is structurally divided into three regions: foregut, midgut, and hindgut, and plays a major role in maintaining this osmotic and ionic balance at benign conditions (MacMillan and Sinclair, 2011a). Mechanical breakdown of food occurs in the foregut, the bulk of digestion and nutrient absorption happens in the midgut, and any water remaining is reabsorbed in the hindgut before the digested bolus passes into the rectum and is excreted (Linser and Dinglasan, 2014; Phillips et al., 1987). Additionally, the hindgut and specialized diverticulae known as Malpighian tubules (analogous to human kidneys) work together to maintain renal function (MacMillan and Sinclair, 2011b; MacMillan et al., 2017; Overgaard et al., 2021; Yerushalmi et al., 2018). Water and ions can move across renal epithelia in two primary ways: transcellularly through aquaporins, ion transporters, or channels, or paracellularly through structures called septate junctions (Izumi and Furuse, 2014; Jonusaite et al., 2016; Jonusaite et al., 2017a; Jonusaite et al., 2017b). These junctions are ladder-like protein complexes located between gut epithelial cells that regulate the passive movement of solutes and water (MacMillan et al., 2017; O’Donnell, 2008; Phillips et al., 1987).

Under optimal environmental conditions, transport and leak rates are balanced such that hemolymph water volume and [Na^+^] remain high, while [K^+^] concentration remains low (D’Silva et al., 2017; Harvey et al., 1983; MacMillan and Sinclair, 2011b; MacMillan et al., 2015b; Overgaard and MacMillan, 2017). At low temperatures, ion and water homeostasis become disrupted when a net leak of Na^+^ and water into the gut lumen occurs, reducing hemolymph volume (MacMillan and Sinclair, 2011b). Recent evidence has suggested that gut epithelial barriers of both *Drosophila melanogaster* and *Locusta migratoria* become disrupted in the cold. This failure of barrier function has been hypothesized to contribute to ion balance disruption in the cold by allowing water and/or solutes to leak down their electrochemical gradients (Andersen et al., 2017b; Brzezinski and MacMillan, 2020; MacMillan et al., 2017). In both locusts and *Drosophila*, this leak was observed *in vivo* using the fluorescently labelled dextran (FITC-dextran, 3-5 kD), a molecule used frequently in epithelial barrier research because it is too large to move transcellularly and cannot be metabolized by animals (Andersen et al., 2017b; Brzezinski and MacMillan, 2020; Jensen-Jarolim et al., 1998; MacMillan et al., 2017; Woting and Blaut, 2018). Brzezinski and MacMillan (2020) found that dextran leak in the cold occurs unidirectionally from the gut to the hemocoel when fed to locusts, but not in the opposite direction when injected into the hemocoel, implying that gut contents may be particularly likely to leak into the hemocoel of insects during cold stress.

In addition to regulating the flow of water and ions, the gut also houses an abundant microbial community which is primarily composed of bacteria and yeasts (Dillon and Dillon, 2004; Padilla, 2016; Wong et al., 2011). Recent studies suggests that gut bacteria and yeasts may affect an insect’s survival at low temperatures. For example, *D. melanogaster* with a healthy gut flora exhibit significantly increased cold tolerance (Henry and Colinet, 2018; Moghadam et al., 2018; Padilla, 2016), However, immune activation, typically associated with bacterial pathogens, has also been reported in adult and larval *D. melanogaster* in response following cold stress. Specifically, the cold stress response is characterized by increased expression of genes in the Toll, immune deficiency (Imd), and/or Janus kinase (JAK)-signal transducer and activator of transcription (STAT) pathways (Salehipour-shirazi et al., 2017; Sinclair et al., 2013; Štětina et al., 2019).

These repeated reports of immune activation in the cold could putatively be explained by the presence of bacteria in the hemolymph that originate from the “leaky” gut, but no studies so far have directly tested if gut bacteria leak into the hemocoel of insects after a cold exposure. Here, we used *in vivo* and *ex vivo* experiments with migratory locusts to test the hypothesis that immune activation, previously observed in cold stressed insects, is a direct response to cold-induced bacterial leak from the gut.

## Materials and methods

### Locust rearing

Locusts (*Locusta migratoria*) used in the experiments were derived from a colony maintained at Carleton University, Ottawa, ON. The colony was reared on a 16 h:8 h light:dark cycle at 28°C at 60% relative humidity, under crowded conditions. All locusts were provided with a dry mixture of oats, wheat germ, wheat bran, and dry milk powder, as well as fresh wheat clippings, three days a week *ad libitum.* All locusts used in experiments were sexed and equal numbers of males and females were used in all experiments.

### Chill-coma recovery time and chilling injury scores

Cold tolerance of locusts was quantified using chill coma recovery time (CCRT) and degree of chilling injuries (injury score). The methodology used here was slightly modified from Brzezinski and MacMillan (2020). Locusts from the colony were collected on the day of the experiment and placed individually in 50 mL polypropylene falcon tubes (*n* = 10 for each group). Holes were made in the lids of the tubes, which provided locusts access to air for the duration of the experiment. Control locusts were placed in an incubator (Isotemp BOD Refrigerated Incubator 3720A; Thermo Fisher Scientific, ON, Canada) with dry oat mixture and fresh wheat clippings for 48 h at 25°C. Locusts undergoing cold stresses were suspended in a circulating cooling bath (Model AP28R-30; VWR International, Radnor, PA, USA) using a Styrofoam rig. The bath was filled with 100% ethylene glycol, pre-set to 25°C, and cooled to −2°C at a rate of - 0.25°C min^-1^. Locusts were then left at −2°C for 12, 24, 36, or 48 h. Temperature was monitored and confirmed throughout the duration of the cold exposure using type-K thermocouples (TC-08 Data Logger; Picotech, Texas, USA) in the glycol and in the hemocoel of an additional locust not used in the experiments. After each time point, locusts, in their comatose state, were removed from their tubes and placed on their side on a clean bench at room temperature (24.0-25.0°C). To measure CCRT, each locust was closely monitored for the time taken to regain neuromuscular function and stand on all six legs, which was recorded as that locust’s CCRT. A 90 min cut-off point was used, after which any locust that had failed to stand on all legs was marked as “unrecovered”.

After 90 min, locusts (regardless of their state) were placed in clean 50 mL polypropylene tubes with dry oat mixture and fresh wheat clippings and left to recover in the incubator at 25°C. After 24 h at 25°C, an injury assessment was done by a single assessor (M.E) using a 5-point scale adapted from MacMillan et al. (2014). Scores were defined as follows: 0: no movement observed (dead); 1: limb movement (leg and/or head twitching); 2: greater limb movement (leg extension and retraction, and whole body twitching), but unable to stand; 3: able to stand, but unable or unwilling to walk or jump; 4: able to stand, walk, and or jump, but lacks coordination; and 5: movement restored similar to pre-exposure levels of coordination. After scoring injury, locusts were returned to their respective tubes with replenished oats and wheat clippings and placed in the incubator. The injury assessment was repeated three more times for each locust – two, three, and four days (48, 72, 96 h) post-cold exposure.

### Development of fluorescent bacteria feeding protocol

To be confident that any bacteria present in the hemolymph originated from the gut lumen, we developed a novel feeding and bacterial leak assay. This protocol used a mutant fluorescent strain of *E. coli, GFPmut3* (λ_max_ excitation: 500 nm, λ_max_ emission: 513 nm) with a green fluorescent gene on a plasmid alongside an ampicillin-resistant gene (Chalova et al., 2008; Zhao et al., 2008) and preliminary trials were conducted to optimize the assay. In the final protocol, an overnight culture of *GFPmut3 E. coli* was grown in LB Broth containing ampicillin (100 μg ampicillin 1 mL^-1^ media) in an incubator at 37°C (Model MIR-154; PHC Corporation, Wood Dale, IL, USA). Then, 125 mL of medium was centrifuged at 8000 rpm (9730 x *g*) for 15 min (Sorvall RC 6 Plus; Thermo Scientific, Waltham, MA, USA). The supernatant was discarded, and the bacterial pellet was resuspended in 1.25 mL of distilled water in a 2 mL centrifuge tube, effectively concentrating the bacterial solution 100-fold. Ten 3 cm strands of freshly cut wheat were added to the tube and allowed to soak for 24 h in an incubator at 37°C. Simultaneously, locusts that were to be used in the experiments were moved to a separate cage for 24 h with no wheat or oats to fast (which ensured they would eat the soaked wheat when presented with it). Each locust was then placed in separate plastic containers (36.3 x 25.1 x 59.9 centimeters), with holes in the lid, along with their own bacteria-soaked wheat where they fed for 24 h in an incubator at 25°C. Further details on the validation of this protocol are included in the supplementary material.

### Investigating cold-induced bacterial leak from the gut

To test whether bacteria in the gut leak into the hemolymph during cold stress, locusts were suspended in a cooling bath at −2°C for 12, 24, 36, or 48 h (*n* = 6, *n* = 10 for 48 h group) following 24 h of feeding on the wheat soaked in the fluorescent bacteria solution. Control locusts were kept in the incubator at 25°C for 48 h with wheat and oats provided *ad libitum.* To collect hemolymph, locusts were pricked dorsally at the head-thorax junction. Hemolymph was then collected using techniques adapted from Findsen et al. (2013). A 50 μL capillary tube was used to collect hemolymph via capillary action at the site of injury. By inserting a pipette tip at the end of the capillary tube, 10 μL of hemolymph was drawn and pipetted into 190 μL of sterile locust saline (in mmol L^-1^: 140 NaCl, 8 KCl, 2.3 CaCl2 Dihydrate, 0.93 MgCl2 Hexahydrate, 1 NaH2PO4, 90 sucrose, 5 glucose, 5 trehalose, 1 proline, 10 HEPES, pH 7.2) in a centrifuge tube, and this process was completed twice to generate two hemolymph samples from each animal. After briefly vortex mixing, one of the 200 μL solutions was spread on a petri dish containing LB with ampicillin, and the other solution was spread on an LB agar plate without ampicillin. Both plates were then incubated at 37°C. The plates were checked for colony growth every day for four days. Colony forming units (CFU μL^-1^) in extracted hemolymph samples were then determined using serial dilution plating on LB agar plates containing ampicillin.

Bacterial leak could plausibly occur following, rather than during, a cold stress, so we performed a follow up experiment to test for bacterial leak following a 6 h rewarming period after the cold stress. Cold exposures were done in an identical manner as described above following bacterial feeding. In this case, however, locusts (*n* = 6 in each group) were placed in small plastic deli containers after the cold stress, with freshly cut wheat (not soaked in the bacterial solution) and dry oat mixture in excess. The containers with the locusts were then placed in the incubator and left to recover at 25°C for 6 h.

To ensure that fluorescent bacteria that are present in the hemolymph of cold-stressed locusts could be recovered using our extraction method, we included a positive control. Locusts (*n* = 4) were suspended in a cooling bath and were left undisturbed while the bath ramped down to −2°C. Individuals were then removed and were injected dorsally at the head-thorax junction with 10 μL of a 1.46 x 10^8^ CFU mL^-1^ solution of *GFPmut3 E. coli* in sterile locust saline. The injection site was then sealed with high vacuum grease (Dow Corning, Etobicoke, ON, Canada) before returning the locusts back to the cooling bath for 1 h at −2°C. Hemolymph samples were then collected as above.

### Determining the role of gut bacteria in cold-induced paracellular barrier disruption

To better investigate cold-induced paracellular leak across the gut *ex vivo* and in the absence of the natural gut microbiome, we tested for leak of FITC-dextran (FD4; 3–5 kDa, Sigma-Aldrich, St Louis, MO, USA) from isolated gut segments of *L. migratoria* using a modified gut sac approach (Gerber and Overgaard, 2018; Hanrahan et al., 1984). We modified the preparation by 1) not everting the gut segments, and 2) inserting two separate pieces of polyethylene tubing on the anterior and posterior side of each segment. On the day of the experiment, locusts were decapitated and prepared for dissection by removing all appendages. The thorax and abdomen were placed in a Sylgaard-lined petri dish and locust saline was used to keep tissues moist during dissections. An incision was made in the anterior to posterior direction along the ventral side to pin open the body cavity. All structures aside from the gut tract were cleared away before the segments were isolated. Portions of the gut were isolated in a manner convenient to the method being applied rather than by anatomical definition. Briefly, the segments were described as follows: anterior, from the anterior-most portion of the esophagus to the midgut caecae, central, from posterior of the midgut caecae to the midgut-hindgut junction where the Malpighian tubules connect, and posterior, from the anterior-most portion of the ileum to the posterior end of the rectum.

To suspend each isolated gut segment within our system, a heat flared polyethylene tube (PE tube; VWR ID x OD: 0.023 x 0.038”, Radnor, USA) was inserted and tied into the anterior margin of the section. Once secure, standard locust saline was injected through the PE tube to thoroughly rinse out the gut contents, including the vast majority of the gut microbiome. A second heat flared PE tube was then inserted and tied into the posterior margin of the segment. Preparations were kept in a petri dish containing continuously oxygenated (95% O2, 5% CO2, Praxair, Danbury, USA) saline at room temperature (23°C) until all three segments had been prepared for suspension. Once complete, a 9.6 x 10^-4^ M FITC-dextran solution was injected via PE tube into each preparation until it had filled both PE tubes, ensuring the lumen was filled with the saline containing the FITC-dextran. Each preparation was suspended in a beaker containing 25 mL of continuously oxygenated locust saline, which acted as our serosal environment. After a 30 min rest period at room temperature to allow for tissue stabilization, the beaker was moved into the cooling bath (preset at 0°C) and monitored for 5 h (Gerber and Overgaard, 2018). Preparations were then removed from the cooling bath and monitored for an additional 2 h at room temperature to account for any effect that rewarming had on the rate of leak.

Throughout the experiment, 90 μL aliquots were collected directly from the beakers once every hour for the duration of the experiment and transferred to a 96-well plate (Corning Falcon Imaging Microplate; black/clear bottom) for fluorescence spectrophotometry (λ_max_ excitation: 485 nm, λ_max_ emission: 528 nm; BioTek Cytation 5 Imaging Reader, Winooski, USA). Concentrations of FITC-dextran in the samples were determined by reference to a standard curve of FITC-dextran in locust saline. The results obtained from fluorescent analyses were then plotted to obtain leak rates per cm^2^ of gut tissue for all preparations. Briefly, the slope ([FITC-dextran] (μmol) against Time (h)) of each gut sac sample and measurements of tissue length and width (treated as cylindrical surface area) were used to calculate leak rates per cm^2^ of gut tissue.

### Data analysis

All collected data were analyzed in R Studio version 3.5.3 (R Core Team, 2019). The distribution and variance of residuals were assessed using Shapiro-Wilk tests and Q-Q plots, which supported the use of non-parametric tests. The effect of cold duration on CCRT was analyzed with a Kruskal-Wallis (KW) test followed by pairwise Wilcoxon tests with the Benjamini-Hochberg (BH) correction. Since the assumption of normality was not met, a generalized linear model (glm) with a Poisson distribution was used to test for the effect of cold duration and assessment day on injury scores. A KW test followed by pairwise Wilcoxon tests (with BH correction) were used to test for significant differences in injury scores on the first and fourth assessment days following the cold exposures. Again, because of the non-normal distribution of the data, a glm with a Poisson distribution was used to examine the effect of bacterial feeding on chilling injuries. Cold exposure duration was held as a fixed categorical variable, while assessment day was held as a continuous variable (in their respective analyses). For the *ex vivo* gut sac experiment, FITC-dextran leak/cm^2^ of tissue in the cold and post-cold was analyzed via the lmer() function (lme4 and lmerTest packages for R). Time was held as a continuous factor, gut segment as a fixed effect, and each individual locust as a random effect. Finally, paired t-tests were done to compare the rates of FITC-dextran leak/cm^2^ of gut tissue in the cold and post-cold. Log10 of the *ex* vivo gut data were used for statistical analyses. Values presented on all graphs are shown as mean ± standard deviation with the α-level being 0.05 for all statistical tests. For the gut bacterial leak assay, no statistics were used to analyze growth on plates due to lack of any colonies observed from the cold-stressed locusts.

## Results and discussion

### Chill-coma recovery time and survival score following cold exposures

Chill-coma recovery times significantly increased from 19.3 ± 7.07 min after 12 h at −2°C to 80.4 ± 2.95 mins after 48 h at −2°C (for those locusts that recovered within the 90 min cut off period; KW, χ^2^ = 17.67, *P* < 0.001). Longer cold exposures also resulted in fewer locusts recovering before the 90 min cut-off. Exposure to −2°C for 12 or 24 h yielded a 100% recovery rate, which decreased to 75% at 36 h and 37.5% at 48 h (Figure 1A).

**Figure 1.**
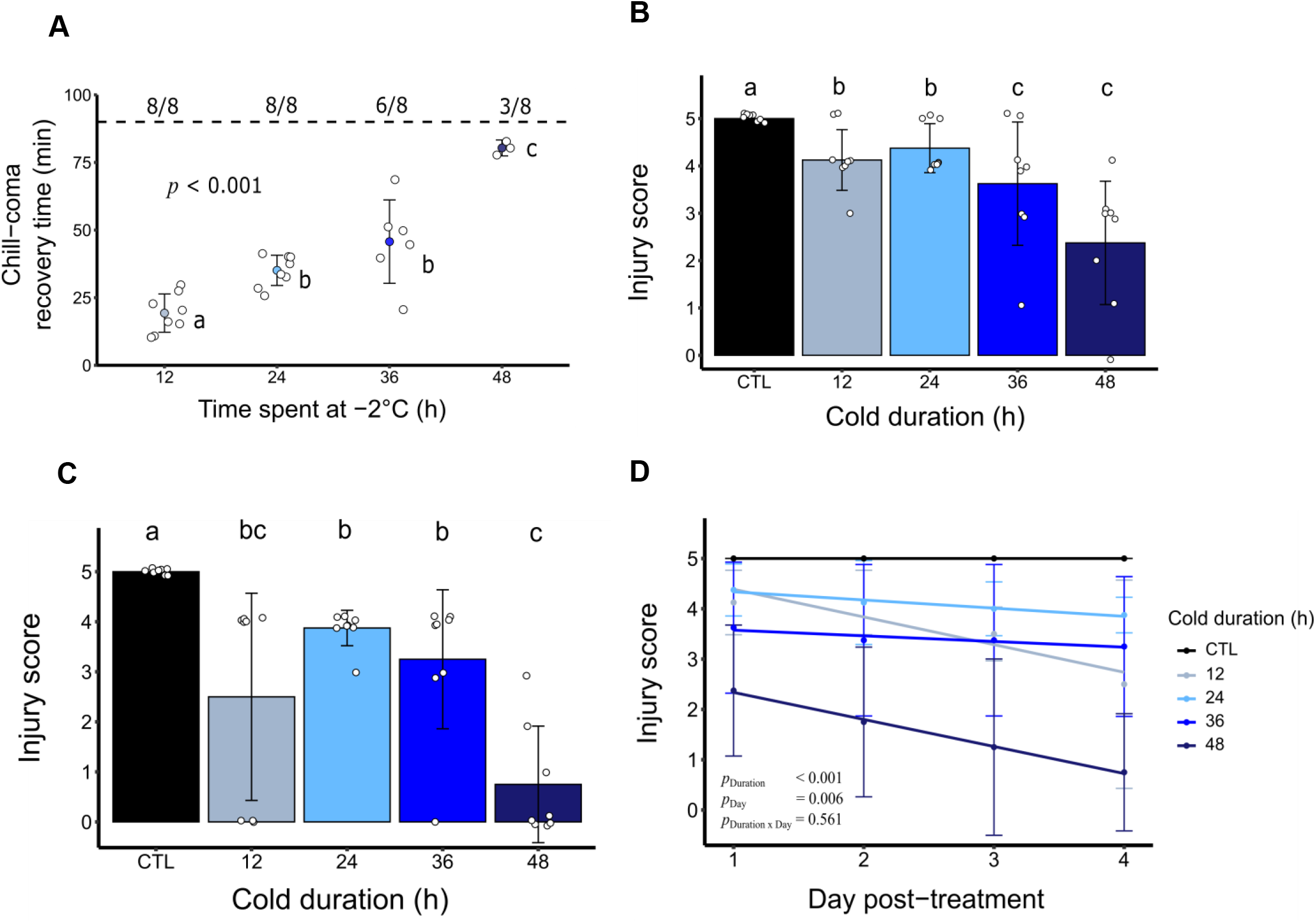
Chilling injuries in *L. migratoria* increase with longer cold stresses.) **A)** Locusts were observed for 90 mins post-chilling and were marked as having recovered when they were able to stand on all six legs. Values above the dashed black line represent the number of locusts in each cold exposure group that recovered before 90 min. Open points represent individual CCRT for each locust.. Locust injury scores were then assessed **B)** one day and **C)** four days after cold exposures (*P* < 0.001 for both plots). Groups sharing the same letter are not significantly different. Open data points represent individual injury scores, and are slightly vertically scattered around scores for clarity. **D)** Plot showing the decline in survival over the four day period for each group (*n* = 8). Data points shown represent mean ± s.d.

Chilling injuries were quantified using injury scores that clearly demonstrated how longer cold exposures led to higher degrees of injury (lower injury scores; GLM, *F_4,140_* = 44.62, *P* < 0.001) (Figure 1B and C). Over the four days following the cold stresses, chilling injuries worsened (GLM, *F_3,140_* = 4.30, *P* = 0.006; Figure 1D). This effect, however, was only significant when we included the 48 h cold exposure group in the statistical model, suggesting that particularly severe injuries get progressively worse after the cold stress (compare Figure 1B and C).

As previously found in the same species (e.g. Andersen et al., 2017a; Brzezinski and MacMillan, 2020; MacMillan et al., 2014) we here find that increasing durations of exposure to low temperature (−2°C) results in higher degrees of injury (Fig. 1). Latent chilling injuries manifested in locusts exposed to −2°C for 48 h, quantified by decreasing injury scores (more severe chilling injuries) over a three day period after the first survival assessment. A similar pattern was found in *D. melanogaster* exposed to a relatively long cold stress (25 h) at 0°C; where lower injury scores one day after the cold stress led to significantly more deaths the following three days after removal from the cold (El-Saadi et al., 2020). These results from *D. melanogaster* and *L. migratoria* suggest that chilling injuries do not fully heal and may even continue to worsen in the days following removal of the insect from a severe cold stress (Figure 4; El-Saadi et al., 2020).

### Bacterial leak across the gut following cold exposures

If the continuous decline in survival is a result of bacterial infection or an adverse immune response (Sadd and Siva-Jothy, 2006), then this could be explained by bacteria leaking from the gut lumen into the hemocoel during or following a cold stress. To test this, we fed locusts a fluorescent strain of *E. coli* before exposing them to the cold. Immediately following a cold exposure, hemolymph samples were taken from locusts and spread on LB agar plates with or without ampicillin to look for *GFPmut3* colony growth exclusively or any colony growth, respectively. No bacterial colonies were seen on any plates (LB agar plates with or without ampicillin) containing hemolymph from the locusts, regardless of cold exposure duration (Table 1). This finding was confirmed in an experiment to examine bacterial leak following a period of rewarming where locusts were left to recover with food at benign temperature for 6 h following the cold exposure. Similar to the acute experiments, no bacterial colonies were observed on any plates regardless of cold exposure duration (Table 1).

**Table 1.** Bacterial CFU after plating locust hemolymph immediately, or 6 h, following cold exposures at −2°C. A positive control group was injected with a bacterial solution before extracting hemolymph after 1 h at −2°C. Male and female locusts in all groups were used in a 1:1 ratio.

| Experimental design | Sample size | Bacterial concentration (CFU mL<sup>-1</sup> hemolymph) |
| --- | --- | --- |
| Positive control | 4 | $5.46 \times 10^5 \pm 8.34 \times 10^4$ |
| Immediately after cold stress |  |  |
| Control | 6 | 0 |
| 12 h | 6 | 0 |
| 24 h | 6 | 0 |
| 36 h | 6 | 0 |
| 48 h | 10 | 0 |
| 6 h following cold stress |  |  |
| Control | 6 | 0 |
| 12 h | 6 | 0 |
| 24 h | 6 | 0 |
| 36 h | 6 | 0 |
| 48 h | 6 | 0 |

In another Orthopteran species, the spring field cricket (*Gryllus veletis*), the ability of the immune system to clear bacteria from the hemolymph is significantly reduced at low temperatures (Ferguson et al., 2016). This is also seen in some other orders of insects (Ferguson and Sinclair, 2017). As hemolymph was plated immediately after the cold stresses in one of our experiments, it is unlikely that the absence of bacteria was a result of immune-related bacterial clearance. In this case, there are two probable scenarios: 1) gut barriers maintain their integrity well enough to prevent septicemia, despite damage to the gut epithelia and leak of solutes, or 2) gut epithelia retain their barrier properties in the cold. From our data (see below) and those of Gerber and Overgaard (2018), the degree of FITC-dextran leak in the cold is insufficient to support a purported increase in gut barrier permeability which would have allowed bacteria to cross over in our experiments. Hence, the second explanation is more likely.

Since the leakage of gut bacteria is unlikely to explain the reported cold induced immune activation following cold stresses, one possibility is that this response is associated with the leak of immunogenic components of bacteria such as lipopolysaccharide (LPS). Although purified LPS has been shown to not activate the Imd pathway in *Drosophila* (Kaneko et al., 2004), it does induce overexpression of antimicrobial peptide genes in silkworms (Tanaka et al., 2009). Another possibility is the gradual development of cellular damage and an associated leak of intracellular proteins such as actin. Tissue injury or cell death can lead to the release of actin into the hemolymph of insects (Dominguez and Holmes, 2011). Srinivasan et al. (2016) clearly show that actin in the hemolymph of *D. melanogaster* elicits an immune response via JAK/STAT pathway activation. Cold-stressed insects exhibit a disrupted cytoskeleton in cells (Cottam et al., 2006; Des Marteaux et al., 2018) and cold exposures lead to an upregulation of genes important in the maintenance of the cytoskeleton (Kim et al., 2006; MacMillan et al., 2016; Teets et al., 2012), providing further support to this hypothesis.

### FITC-dextran leak across epithelia of isolated gut segments

To examine whether the locust gut becomes leaky enough in the cold to permit increased FITC-dextran movement, we used a modified gut sac technique to examine *ex vivo* leak of FITC-dextran in the absence of the vast majority of the natural gut microbiota. At 0°C, no significant differences between FITC-dextran leak rates were found between the segments (Fig. 2; LME, *F_2, 15_* = 1.03, *P* = 0.379). The same was true for segments held at room temperature over the course of experiments (LME, *F_2, 10_* = 0.989, *P* = 0.406). When comparing between both temperature treatments (−2°C or 23°C), FITC-dextran leak rates did not significantly differ between segments (LME, *F_2, 30_* = 0.882, *P* = 0.424) or between treatments (LME, *F_1, 30_* = 1.04, *P* = 0.315). Furthermore, there was no statistically significant interaction between the gut segment and type of treatment received when analyzing FITC-dextran leak rates (LME, *F_2, 30_* = 1.10, *P* = 0.346; Figure 2). Finally, no significant differences were found between leak rates after 5 h of cold stress and after 2 h at 23°C following cold stress in any of the three segments (foregut: two-tailed *t*_5_ = 0.668, *P* = 0.534; midgut: two-tailed *t*_5_ = 1.19, *P* = 0.288; hindgut: two-tailed *t*_5_ = 1.27, *P* = 0.261).

**Figure 2.**
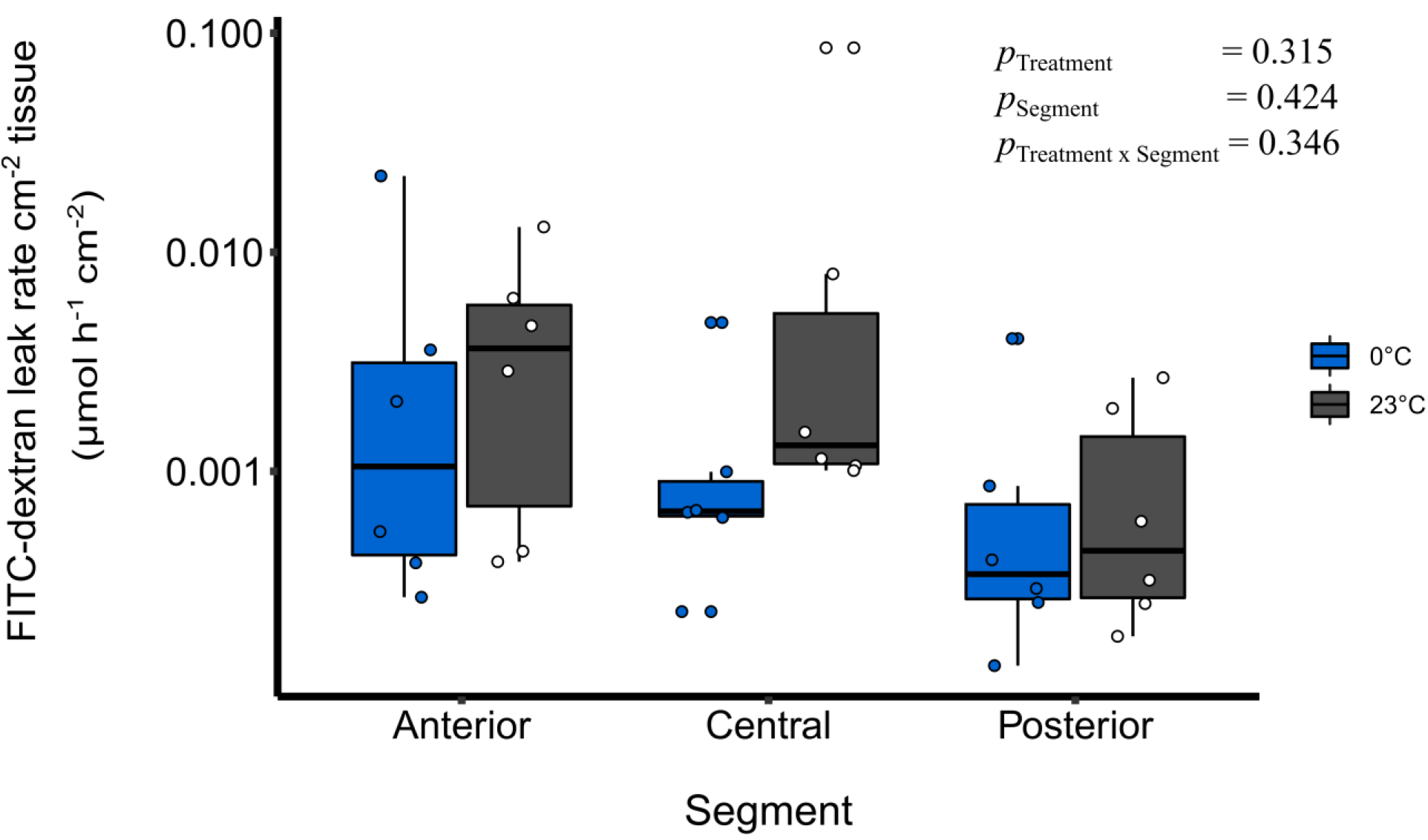
Leak rates of FITC-dextran per cm^2^ of tissue are similar both in the cold and at room temperature. Log10 values of FITC-dextran leak rates are shown foreach of the three segments (anterior, central, and posterior; n=6 samples per treatment, with n=12 overall per segment). Box plot midlines represent median values. Blue and white-filled points represent individual samples taken per treatment.

While the underlying mechanisms of cold-induced immune activation remain unclear, we also discuss to what extent the insect gut becomes leaky at low temperatures. When we exposed isolated gut segments to the cold *ex vivo*, we found no significant increase in the rate of FITC-dextran leak from the lumen to the surrounding solution compared to isolated gut sacs at benign temperature (Figure 3). This agrees with a previous study on the same species using everted rectal sacs *ex vivo,* where no change was found in the mucosal-to-serosal clearance of FITC-dextran in the cold (Gerber and Overgaard, 2018). Together, these data now lead us to question not only the validity of FITC-dextran as a marker of paracellular permeability, but also whether gut epithelial barriers become leaky at all. A common factor in the studies that reported FITC-dextran leak in the mucosal to serosal direction at low temperatures was that the FITC-dextran was administered to the insects orally. The ingestion of FITC-dextran means that it would be concentrated in the gut initially, where there also exists an abundant microbial community. Some bacterial species produce dextranases which cleave larger dextran molecules into smaller fragments (Khalikova et al., 2005). If these bacterial species are present in the gut of healthy locusts, then it would explain the leak of FITC-dextran at benign temperature which would most likely arise from the fragmentation of the large FITC-dextran molecule into smaller polysaccharides.

The findings from the present study may shift our understanding of barrier failure in cold stressed insects and raise questions as to what may trigger an insect’s immune system in the cold. Considering that we did not find any evidence of bacterial leak from the gut, we propose that immune activation in the cold may arise from sterile causes, possibly as a result of cell damage from chilling injuries, or immunogenic molecules from microbes. From this standpoint, we suggest future studies should attempt to examine cold-induced immune activation by looking at damage- or pathogen-associated molecular patterns such as actin and lipopolysaccharide in the hemocoel following cold stresses.

## Acknowledgements

We wish to thank Jeffery Dawson and Marshall Ritchie for assistance with locust rearing and colony maintenance; Amanda Carroll for help with procuring bacterial stocks and early troubleshooting with microbiological techniques, and Laura Ferguson for valuable discussion in the early stages of the study’s design.

## Author contributions

Conceptualization: M.E., H.M., L.G.; Methodology: M.E., H.M., A.W., A.H., L.P., L.G., J.O; Resources: H.M., A.W.; Data curation and analysis: M.E., K.B.; Writing – original draft: M.E.; Writing – review and editing: H.M., J.O., A.W., K.B., L.G., A.H., L.P., M.E.; Visualization: M.E.; Supervision: H.M.; Project administration: H.M.; Project funding: H.M.

## Data availability

All data is provided as a supplementary file for review.

